# Diminished miRNA activity is associated with aberrant cytoplasmic intron retention in ALS pathogenesis

**DOI:** 10.1101/2021.01.27.428555

**Authors:** Marija Petric-Howe, Hamish Crerar, Jacob Neeves, Giulia E. Tyzack, Rickie Patani, Raphaëlle Luisier

## Abstract

Intron retention (IR) is now recognized as a dominant splicing event during motor neuron (MN) development, however the role and regulation of intron-retaining transcripts (IRTs) localized to the cytoplasm remain particularly understudied. By resolving the spatiotemporal dynamics of IR underlying distinct stages of MN lineage restriction, we identify a cytoplasmic group of IRTs that is not associated with reduced expression of their own genes but instead with an upregulation of predicted target genes of specific miRNAs, the motifs of which are enriched within the intronic sequences of this group. Next, we show that ALS-causing VCP mutations lead to a selective increase in IR of this particular class of introns. This in turn temporally coincides with an increase in the expression level of predicted target genes of these miRNAs, providing a potential mechanistic insight into ALS pathogenesis. Altogether, we propose a novel role for the cytoplasmic intronic sequences in regulating miRNA activity through miRNA sequestration, which potentially contributes to ALS pathogenesis.

## INTRODUCTION

Intron retention, a mode of alternative splicing whereby one or more introns are retained within a mature polyadenylated mRNA, has been greatly understudied in mammalian systems and for a long time mostly considered as a product of inefficient or mis-splicing. With advances in detection strategies, IR became recognised as a more widespread and regulated process than previously thought, and the idea that IR could even functionally modulate cellular processes has come into focus, with its role(s) in cellular physiology beginning to unfold (1, 2).

Neural cells exhibit a higher proportion of retained introns compared to other cell types and there is an expanding body of evidence demonstrating a functional role for intron retention (IR) both in neuronal development and homeostasis (1, 3–5). Transcripts that exhibit IR often remain in the nucleus, mostly considered to be a means of reducing the expression levels of transcripts not required for cellular physiology at a particular stage (2, 6, 7). Some of these transcripts would eventually be degraded by the nuclear exosome, while specific signals could stimulate splicing of the retained intron in others, resulting in export of the fully spliced mRNA into the cytoplasm and its subsequent translation (5). Indeed, nuclear detention of intron-retaining transcripts (IRTs) provides a powerful mechanism to hold gene expression in a suppressed but poised state that allows rapid protein production if and when an appropriate stimulus is received (4, 5, 8–10).

Although the stable cytoplasmic localisation of intronic sequences in neurons has been reported since 2013 (11), there has been limited investigation into the possible role of cytoplasmic IRTs. This has presumably been overlooked in part due to detection limitations, but also due to a notion that these transcripts would likely contain premature translation termination codons (PTCs) and as such, be degraded by nonsense mediated mRNA decay (NMD) (12). Whilst examples of IR coupled with NMD have been found to downregulate gene expression, such as in granulocyte development (13), these transcripts can encounter other fates in the cell (14). Indeed, one of the few studies focussing on cytoplasmic IR in neurons showed an ‘addressing’ function for intronic RNA sequences, determining the spatial localization of their host transcripts within cellular compartments such as dendrites (15). Another speculated function of IR has arisen following the identification of miRNA binding motifs within the retained intronic sequences. This offers an intriguing route through which miRNA-directed degradation pathways might regulate abundance of IRTs; alternatively, the retained introns themselves may serve as miRNA sinks, or even encode novel miRNAs termed mirtrons (16, 17). Altogether, despite advances in our understanding of IR in neuronal cells, much remains unanswered.

The importance of investigating the roles of IR has been further corroborated by studies that demonstrate its relevance across a diverse range of neurodegenerative diseases (18–21). One such example is amyotrophic lateral sclerosis (ALS) (20), a rapidly progressive and incurable disease, which leads to selective degeneration of motor neurons (MNs). ALS is characterised by protein inclusions and axonal degeneration, and is often associated with RNA processing defects. ALS-causing mutations occur in numerous genes encoding crucial regulators of RNA-processing, which are normally expressed throughout development. Despite the growing number of causative gene mutations being identified in ALS, the precise aetiology remains unknown and early molecular pathogenic events remain poorly understood. We previously made the novel discovery that aberrant IR is a widespread phenomenon in ALS (20), which was corroborated by subsequent studies (21, 22). Moreover, we went on to demonstrate aberrant cytoplasmic IR as a widespread molecular phenomenon in VCP-related ALS (23). We showed that ALS-related aberrant cytoplasmic IRTs have conspicuously high affinity for RNA binding proteins (RBPs), including those that are mislocalized in ALS and proposed that a subset of cytoplasmic intronic sequences serve as ‘blueprints’ for the hallmark protein mislocalization events in ALS (24, 25). This raises an exciting possibility that intronic RNA sequences play additional significant roles beyond their recognized nuclear function. Nevertheless, the role and physiological relevance of cytoplasmic IR during neuronal development and disease still remains largely unresolved.

Against this background we sought to characterise the spatiotemporal dynamics of IRTs by re-analysing RNA-seq data from nuclear and cytoplasmic fractions of patient-specific hiPSCs undergoing motor neurogenesis. We first show that retained introns exhibit compartment-specific features including their dynamics, biological pathways, and molecular characteristics during this process. We reveal a specific class of retained introns in the cytoplasm that is not associated with gene expression changes but exhibits high miRNA binding potential, which is functionally validated by identifying an altered expression profile of the predicted miRNA target genes. We finally analyze this class of retained introns in stem cell models of familial ALS and find evidence for a functional depletion of specific miRNAs, possibly as a result of cytoplasmic intronic sequences-mediated sequestration, which has potential implications for ALS pathogenesis and the development of therapies in this devastating and incurable disease.

## MATERIALS AND METHODS

### Compliance with ethical standards

Informed consent was obtained from all patients and healthy controls in this study. Experimental protocols were all carried out according to approved regulations and guidelines by UCLH’s National Hospital for Neurology and Neurosurgery and UCL’s Institute of Neurology joint research ethics committee (09/0272).

### RNA-sequencing data

We obtained paired-end polyA stranded RNAseq libraries prepared from fractionated nucleus and cytoplasm obtained from 6 distinct stages of motor neuron differentiation from control and VCP^*mu*^ samples (iPSC, and days 3, 7, 14, 21 and 35; **Supplementary Table S1**) from previously published study (**GSE152983**) (23). We also obtained paired-end RNA sequencing reads derived from one independent study on familial form of ALS caused by mutant SOD1 (n=5; 2 patient-derived SOD1A4V and 3 isogenic control MN samples where the mutation has been corrected; Hb9 FACS purified MNs, **GSE54409** (26).

### Transcript and Gene expression analysis

Kallisto (27) was used to (1) build a transcript index from the Gencode hg38 release Homo sapiens transcriptome (-k 31), (2) pseudo-align the RNA-seq reads to the transcriptome and (3) quantify transcript abundances (-b 100 -s 50—rf-stranded). Subsequent analysis was performed with the R statistical package version 3.3.1 (2016) and Bioconductor libraries version 3.3 (R Core Team. R: A Language and Environment for Statistical Computing. Vienna, Austria: R Foundation for Statistical Computing; 2013). Kallisto outputs transcript abundance, and thus we calculated the abundance of genes by summing up the estimated raw count of the constituent isoforms to obtain a single value per gene. For a given sample, the histogram of log2 gene count is generally bimodal, with the modes corresponding to non-expressed and expressed genes. Reliably expressed genes/transcripts for each condition (VCP^*mu*^ or control at days 0, 3, 7, 14, 22 and 35 in each fraction) were next identified by fitting a two-component Gaussian mixture to the log2 estimated count gene/transcript data with R package mclust (28); a pseudocount of 1 was added before log2 transformation. A gene/transcript was considered to be reliably expressed in a given condition if the probability of it belonging to the non-expressed class was under 1% in each sample belonging to the condition. 18,834 genes and 102,047 transcripts were selected based on their detected expression in at least one of the 24 conditions (i.e. 6 different timepoints of lineage restriction for control and VCP^*mu*^ in nuclear and cytoplasm). Next we quantile normalized the columns of the count matrices with R package limma (29). For differential gene expression analysis we ran Sleuth (30).

### Splicing analysis

The identification of all classes of alternative splicing (AS) events in motor neuron differentiation was performed with the Vertebrate Alternative Splicing and Transcription Tools (VAST-TOOLS) toolset, which works synergistically with the VastDB web server, a collection of species-specific alternative splicing library files (31). Paired-end stranded RNA-seq reads were first aligned with VAST-TOOLS against the Homo sapiens hg38, Hs2 assembly from VastDB with the scaffold annotation Ensembl v88. This contains 74030 exon skipping events, 153119 intron retention events, 474 microexon events, 20812 alternative 3’ UTR events, and 15804 alternative 5’ UTR events. We then merged files from identical samples but different lanes together and then performed differential splicing analysis over time either for the control or for the VCP mutant samples separately using the vast-tools diff command which takes into account the different biological replicates. We then imported the result tables into R. For an AS event to be considered differentially regulated between two conditions, we required a minimum average ΔPSI (between the paired replicates) of at least 15% and that the transcript targeted by the splicing event in question to be reliably expressed in all samples from the conditions compared i.e enough read coverage in all samples of interest.

### Gene ontology enrichment analysis

GO enrichment analysis was performed using classic Fisher test with topGO Bioconductor package (32). Only GO terms containing at least 10 annotated genes were considered. A p-value of 0.05 was used as the level of significance. On the figures, top significant GO terms were manually selected by removing redundant GO terms and terms which contain fewer than 5 significant genes.

### Mapping and analysis of CLIP data

To identify RBPs that bind to retained introns, we examined iCLIP data for 21 RBPs (33), and eCLIP data from K562 and HepG2 cells for 112 RBPs available from ENCODE (34, 35). Before mapping the reads, adapter sequences were removed using Cutadapt v1.9.dev1 and reads shorter than 18 nucleotides were dropped from the analysis. Reads were mapped with STAR v2.4.0i (36) to UCSC hg19/GRCh37 genome assembly. The results were lifted to hg38 using liftOver (37). To quantify binding to individual loci, only uniquely mapping reads were used.

### Analysis of cis-acting features

MaxEntScan (38) was used to calculate maximum entropy scores for 9-bp 5’ splice sites and 23-bp 3’ splice sites. Intron lengths and GC content were calculated using the hg38 human genome assembly. The intronic enrichment for RBP binding site was obtained by computing the proportion of crosslink events mapping to retained intron compared to non-retained introns of the same genes, accounting for intron length. These were defined in relation to the acceptor and donor splice sites, namely the last 30 nucleotides (nts) of exonic sequence upstream of the 5’ splice site (R1), the first 30nts of intronic sequence downstream of the 5’ splice site (R2), the 30nts in the middle of the intron (R3), the last 30nts of intronic sequence upstream of the 3’ splice site (R4), and the first 30ntsof exonic sequence downstream of the 3’ splice site (R5). These regions were defined based on the past studies of the Nova RNA splicing map (39), which has been determined by the positioning of conserved YCAY clusters as well as by the binding sites identified by HITS-CLIP as reported in (40). The nucleotide-level evolutionary phastCons scores for multiple alignments of 99 vertebrate genomes to the human genome were obtained from UCSC (41, 42) and a median score was derived for each individual intron and defined regions of interest. The RBP crosslink event enrichment scores in each region of interest or in each group of intron was obtained by dividing the fractions of introns in a given group over the fraction in the full list of introns that exhibit at least one crosslink event for a given RBP in the defined region or across the full intronic region.

### Spatiotemporal taxonomisation of the retained introns

We performed singular value decomposition (SVD) on the PIR cytoplasmic versus nuclear values of 94,457 introns in n = 48 cytoplasmic samples and n=47 nuclear samples across the 5 distinct stages of motor neuron differentiation from healthy controls and VCP mutants. We analysed 94,457 introns out of the 153,119 annotated introns in VAST-TOOLS given their overlap with reliably expressed genes. We then selected the components maximally capturing variance in PIR. To visualize the right singular vectors 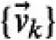, we plotted the PIR on the vertical axis as a function of the time corresponding to each sample on the horizontal axis and coloring all samples corresponding to healthy controls with filled circles, and those corresponding to VCP-mutants in empty circles. Next we identified introns whose PIR profiles correlated (Pearson correlation between individual intron PIR profile and right singular vectors) and contributed (projection of each individual intron PIR profile onto right singular vectors) most strongly (either positively or negatively) with the profile of the singular vectors. In order to identify representative introns for each singular vector, events were ranked according to both projection and correlation scores. The highest (most positive scores in both projection and correlation) and lowest (most negative scores in both correlation and projection) motifs were selected for each singular vector using K-mean clustering.

### MiRNA expression analysis

Total RNA including small RNAs was extracted from “patterned” precursor motor neurons of five control and four mutant cell lines using mirVana™ miRNA Isolation Kit (ambion, life technologies). RNA quantification and its 260/280 ratio were assessed using the nanodrop. Poly(A) tailing and reverse transcription of mature miRNAs was performed using miRCURY LNA RT kit (QIAGEN), with 20 ng of total RNA as input. Reverse transcribed cDNA was quantified using miRCURY LNA SYBR green dye, specifically designed primers (appropriate miRCURY LNA miRNA PCR assays) and QuantStudio 6 Flex Real-Time PCR System (Applied Biosystems). Relative miRNA expression levels between control and mutant cells were quantified using ΔΔCT Method, with U6 snRNA as a reference gene for normalisation. Data was plotted using RStudio software.

## RESULTS

### Nuclear and cytoplasmic IR affect two functionally divergent mRNA subsets

We previously reported a transient IR programme early during human motor neurogenesis using whole-cell RNA-sequencing data (20). To further examine the spatiotemporal dynamics of IR during this process in healthy cells, we re-analysed high-throughput poly(A) RNA-seq data derived from nuclear and cytoplasmic fractions of human induced pluripotent stem cells (hiPSCs; day 0), neural precursors (NPCs; day 3 and day 7), ‘patterned’ ventral spinal motor neuron precursors (pMNs; day 14), post-mitotic but electrophysiologically immature motor neurons (MNs; day 22), and electrophysiologically active MNs (mMNs; day 35) (**Fig. 1A** and **Supplementary Table 1;** 47 nuclear and cytoplasmic samples from 6 time-points; 4 clones from 4 different healthy controls) (23, 43). Using the RNA-seq pipeline VAST-TOOLS (31), we identified 4,189 nuclear and 1,542 cytoplasmic significant alternative splicing (AS) events over time (**Fig. 1B** and **Supplementary Fig. 1A**). In line with our previous study (20), IR was the predominant mode of splicing during neurodevelopment, accounting for 64% and 49% of the included AS events in the nucleus (638 events) and the cytoplasm (541 events) respectively, indicating that cytoplasmic IRTs are more abundant than previously recognized (**Fig. 1B**). Further examining the distributions of percent intron retention (PIR) during MN differentiation in the nucleus and the cytoplasm for 211,501 events revealed that IR exhibits distinct dynamics in the two compartments (**Fig. 1C**). In particular, the nuclear compartment exhibits the highest level of PIR at the hiPSC stage, while the cytoplasmic compartment exhibits the highest level of PIR at DIV=14, which is reminiscent of the early wave of IR we previously reported (20). Notably the cytoplasmic increase in PIR early during differentiation is likely explained by a change in the subcellular localisation of (some) IRTs rather than a modulation of the splicing given the coincident stable level of PIR in the nucleus. Genes related to RNA processing and splicing are among the most affected by IR (13, 44–48) and we previously showed that IR early during MN development specifically affects RNA processing related biological pathways (20). Here we find that cytoplasmic (but not nuclear) IR affects essential genes concerned with mRNA metabolism. In contrast we find that genes targeted by alternative exons (AltEx) are enriched in similar biological pathways in the nucleus and the cytoplasm as shown by Gene Ontology (GO) function analysis (**Fig. 1D**). These findings indicate that the previously reported wave of IR during MN differentiation, using whole-cell RNA-sequencing, likely reflected signals from cytoplasmic IRTs.

**Figure 1.**
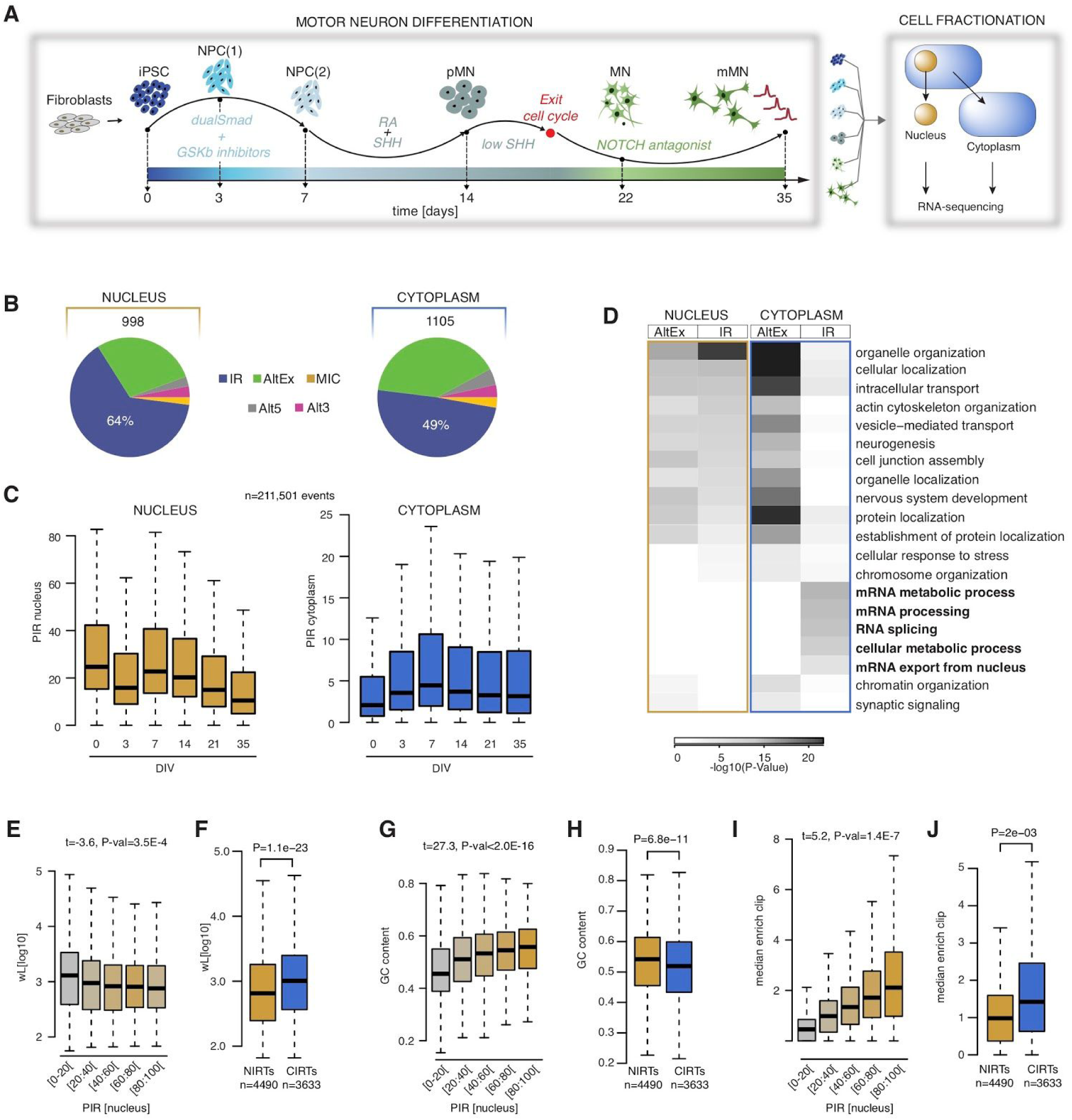
Nuclear and cytoplasmic IR affect two distinct mRNA subsets. **A**. Schematic depicting the iPSC differentiation strategy for motor neurogenesis. Arrows indicate sampling time-points in days when cells were fractionated into nuclear and cytoplasmic compartments prior to (polyA) RNA-sequencing. Four iPSC lines were obtained from four different healthy controls. Induced-pluripotent stem cells (iPSC); neural precursors (NPC); “patterned” precursor motor neurons (ventral spinal cord; pMN); post-mitotic but electrophysiologically inactive motor neurons (MN); electrophysiologically active MNs (mMN). **B**. Pie charts representing proportions of included splicing events at defined stages during motor neurogenesis in nuclear (*left*) and cytoplasmic (*right*) fractions. Total number of events are indicated above the charts. Intron retention (IR); alternative exon (AltEx); microexons (MIC); alternative 5′ and 3′ UTR (Alt5 and Alt3). **C**. Comparison of the percent intron retention (PIR) during MN differentiation in nucleus (*left*) and cytoplasm (*right*) for 21,161 events that exhibit >10% PIR in at least 3 out of 47 nuclear samples. **D**. Heatmap of the GO biological functions enriched among the genes targeted by AltEx or IR in either the nucleus or the cytoplasm. P-values obtained by Fisher enrichment test. **E**. Analysis of the relationship between the PIR in the nucleus and the intron length. Retained introns are grouped in five categories of increasing level of retention in the nucleus as indicated on the *x*-axis. *P*-values obtained from analysis of variance comparing the full model of the logit of maximum IR across all nuclear samples according to the five characteristics with the reduced model removing the characteristic of interest. **F**. Comparison of intron length between nuclear and cytoplasmic retained introns. Nuclear retained introns are defined as intron exhibiting >20% IR in nuclear fraction and <5% IR in cytoplasmic fraction. Cytoplasmic retained introns are defined as intron exhibiting >20% IR in nuclear fraction and >15% IR in cytoplasmic fraction. P-values obtained from Mann-Withney test. **G**. Analysis of the relationship between the PIR in the nucleus and the GC content in %. Data shown as in (**E). H**. Comparison of GC content (%) between nuclear and cytoplasmic retained introns. **I**. Analysis of the relationship between the PIR in the nucleus and the median enrichment for RBP binding site compared to the non-retained introns of the same gene. Data shown as in (**E). J**. Comparison of median enrichment for RBP binding sites between nuclear and cytoplasmic retained introns. For C, E-J-J: Data shown as box plots in which the centre line is the median, limits are the interquartile range and whiskers are the minimum and maximum.

Prior studies reported specific features associated with retained introns including higher GC content, lower intron length and enrichment in RBP binding motifs compared to non-retained introns (1, 47, 49, 50). In line with these studies, we find that the PIR negatively correlates with the intron length and positively correlates with the GC content and the enrichment in crosslink events for 131 RBPs for which CLIP data were available (33–35) both in nuclear and cytoplasmic compartments (**Figs. 1E, G, I** and **Supplementary Fig. 1F**). Additionally we find that i) retained introns are detected in genes containing fewer introns, and ii) the PIR positively correlates with the intronic sequence conservation score (**Supplementary Figs. 1B, D, F**). Surprisingly, however, by specifically comparing the cytoplasmic retained introns (PIR_*NUCLEUS*_ > 20% and PIR_*CYTOPLASM*_ > 15%; **Supplementary Table S2**) with the nuclearly detained retained introns (PIR_*NUCLEUS*_ > 20% and PIR_*CYTOPLASM*_ < 5%; **Supplementary Table S3**) we find that the retained introns that localise to the cytoplasm are on average longer, have lower GC content compared to their nuclear counterparts, have higher RBPs enrichment scores and are more evolutionarily conserved (**Figs. 1F, H, J** and **Supplementary Figs. 1C,E**). Altogether these results show that nuclear and cytoplasmic retained introns exhibit distinct features including their dynamics during human motor neurogenesis, their associated biological pathways, and their molecular characteristics.

### A spatiotemporal taxonomy reveals cytoplasmic retained introns with distinct RBP binding profiles

Having established that nuclear and cytoplasmic IR affect two functionally divergent mRNA subsets, we next used singular value decomposition (SVD) analysis to categorize 94,457 analysed introns into nine groups based on their PIR spatiotemporal dynamics during MN differentiation (**Fig. 2A** and **Supplementary Tables S4-S12**). Of these, three categories are nuclearly-detained retained introns (termed N1-N3 hereafter) and the other six are cytoplasmic retained introns (termed C1-C6 hereafter), that exhibit the following compartment-specific PIR dynamics: stable nuclear (>15%) and low cytoplasmic (<10%) PIR over time (N1), steady reduction in the nuclear PIR over time and stable low cytoplasmic PIR (N2), a transient increase in nuclear PIR and stable low cytoplasmic PIR (N3), steady reduction in both nuclear and cytoplasmic PIR over time (C1), steady increase in nuclear and cytoplasmic PIR over time (C2), increase in cytoplasmic PIR only in terminal differentiation (C3), consistently high nuclear and cytoplasmic PIR over time (C4), early transient increase in nuclear and cytoplasmic PIR (C5) and a late transient increase in nuclear and cytoplasmic PIR (C6).

**Figure 2.**
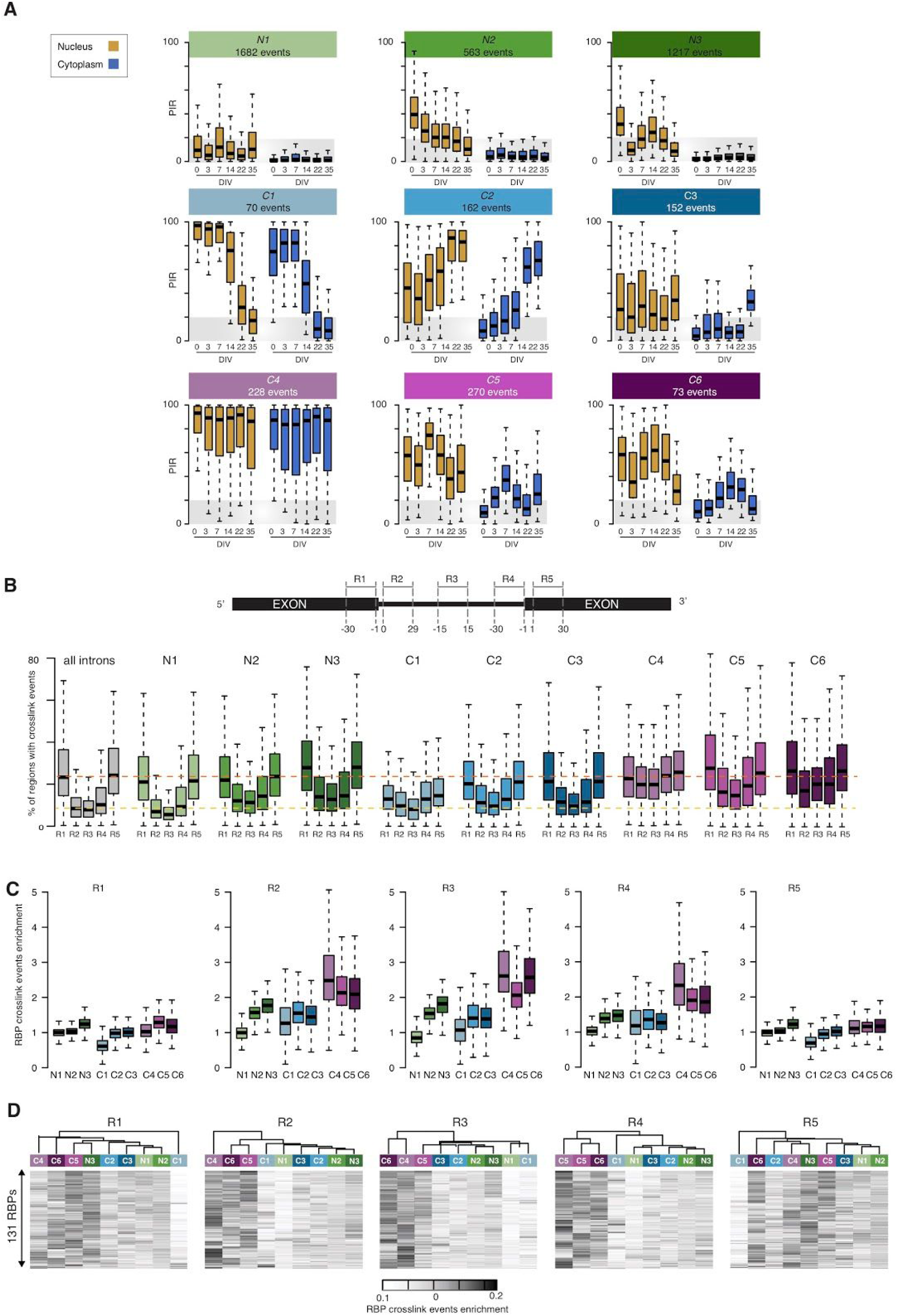
A spatiotemporal taxonomy reveals cytoplasmic IRTs with distinct RBP binding profiles. **A**. Comparison of the nuclear and cytoplasmic percent intron retention (PIR) distributions for 9 groups of retained introns exhibiting distinct spatio-temporal dynamics during MN differentiation as identified using SVD (see Materials and Methods). N1, N2 and N3 contain introns primarily retained in the nuclear compartment while the remaining 6 groups contain introns with significant detection in the cytoplasm. Gold boxes = nucleus; blue boxes = cytoplasm. Grey area indicates the range of PIR values for which an intron is considered non-retained. **B**. (*Upper)* Schematic depicting the selected splicing regulatory regions juxtaposing the splice sites, namely the last 30 nucleotides (nts) of the upstream exon (R1), the first 30nts of 5’ intron region (R2), 30nts in the middle of the intron (R3), the last 30nts of 3’ intron region (R4), and the first 30nts of downstream exon (R5). (*Lower*) Distributions of the percentage of regions in each group of introns that are mapped by at least one crosslink event for each of the available 131 RBPs. **C**. Distribution of the enrichments in crosslink events in each of the selected regions R1, R2, R3, R4 and R5 for the available 131 RBPs across the 9 categories of introns. Enrichment is obtained by dividing the fraction of regions from the group of interest with at least one crosslink event with the fraction of regions from the complete set of introns (n=61872) with a crosslink event. **D**. Heatmaps of the enrichment scores of the crosslinking events for 131 RBPs in the R1, R2, R3, R4 and R5 regions for the 9 groups of introns hierarchically clustered using Manhattan distance and Ward clustering. Data shown as box plots in which the centre line is the median, limits are the interquartile range and whiskers are the minimum and maximum.

Next looking at the percentage of introns per genes targeted by IR in each identified group revealed that IR during MN differentiation does not occur stochastically, but appears to target a specific set of introns in each gene (**Supplementary Fig. 2A**), indicating that additional layer(s) of specific regulation must underlie the regulation of these 9 distinct IR programmes as previously suggested (47). Previous studies suggest that cis-regulatory elements bound by trans-acting factors such as RBPs are likely to play a crucial role in regulating IR (50–52). Thus we next sought to test whether the 9 spatiotemporally distinct classes of IR we identified are associated with different combinations of trans-acting factors that could regulate them. We achieved this by using the publicly available CLIP data to evaluate the crosslink events for 131 RBPs mapping to 5 regions we defined in relation to the acceptor and donor splice sites, namely the last 30 nucleotides (nts) of exonic sequence upstream of the 5’ splice site (R1), the first 30nts of intronic sequence downstream of the 5’ splice site (R2), the 30nts in the middle of the intron (R3), the last 30nts of intronic sequence upstream of the 3’ splice site (R4), and the first 30nts of exonic sequence downstream of the 3’ splice site (R5) (**Fig. 2B**, *upper* and **Supplementary Tables S12-S17**). First, looking at the fractions of regions which are mapped by at least one crosslink event for each RBP, we find that the R1 and R5 exonic regions, sequences of which are the most evolutionarily conserved (**Supplementary Fig. 2B**), exhibit the highest frequency in crosslink events across the 131 RBPs irrespective of the IR grouping, with the exception of the C1 group (**Fig. 2B**, *lower*). These results, which are in line with previous studies showing that the splicing machinery is more likely to form across the exonic regions than across the introns for similarly long introns (>250 nts), indicate that the 9 groups of introns bear similar chances of splicing complex formation with respect to their R1 and R5 exonic regions (53, 54). This is further supported by the finding that identically optimal splicing signals are detected among the 9 groups of introns, with the exception of the C4 group (**Supplementary Fig. 2C**) as failure in splice site recognition (50–52) or decreased expression levels of splicing factors (47) have also been proposed to underlie IR.

Although RBPs exhibit similarly high frequency of binding to the R1 and R5 exonic regions across the majority of the 9 groups of introns, we indeed noticed that the R2, R3 and R4 intronic regions display large variability in the percentages of crosslink events across the different spatiotemporal IR dynamics (**Fig. 2B**, *lower*). Next, looking at the enrichment of RBPs mapping to each of these regions further revealed that the C3, C4 and C5 groups of cytoplasmic retained introns is indeed specifically enriched in RBPs binding to the R2, R3 and R4 intronic regions as opposed to the R1 and R5 regions which display as much RBP binding as the full set of introns (**Fig. 2C**). One of the most enriched RBPs in the R2, R3 and R4 intronic regions is UPF1, an RNA helicase required for nonsense-mediated mRNA decay (NMD) in eukaryotes. UPF1 exhibits high binding occurrence across the five regions of interest for the {C4, C5, C6} groups as opposed to the other groups for which UPF1 is strictly enriched in R1 and R5 regions (**Supplementary Fig. 2D**). Different combinations of RBPs have been shown to coordinately regulate functionally coherent “networks” of exons and introns (55). Thus, using unsupervised hierarchical clustering of the 9 groups of introns based on their 131 RBP enrichment scores profiles we finally showed that {C3, C4, C5} form a coherent regulated group of introns in respect to all selected regions except the R1 exonic region (**Fig. 2D**). Altogether, this analysis identifies a coherent regulated supergroup of retained introns which exhibits specific elements in their intronic regions that are not necessarily related to splicing efficiency, but rather may perform an additional role in the regulation of mRNA metabolism.

### Identification of a cytoplasmic group of IRTs with a high capacity for miRNA sequestration

The finding that a coherent regulated supergroup of retained introns bears similar chances of splicing complex formation with respect to their R1 and R5 exonic regions but with a potential regulatory role in mRNA metabolism through RBP intronic sequence binding, prompted us to look at the occurrence of RBP crosslink events across the entire intronic region as opposed to the 5 predefined regions of interest. We showed that while the {C4, C5, C6} groups have a somewhat higher prevalence of crosslink events (**Fig. 3A**), the C5 group specifically has the highest fraction of introns with at least one crosslink event across the 131 studied RBPs compared to the full set of introns (**Fig. 3B**). Notably, the {C4, C5, C6} groups overall have a lower intron length compared to {C1, C2, C3}, and thus this result cannot be due to a bias in size. Of the 51 RBPs that exhibit >19% higher fraction of binding to the C5 group compared to the full set (Fisher enrichment P-value < 0.01; **Fig. 3C**) at least 9 are key regulators of mRNA transport such as UPF1 and IGF2BP1 (56–58), and 6 RBPs are involved in miRNA regulatory pathways such as DROSHA and PUM2 (59–61), UPF1 being involved in the two (62–64) (**Figs. 3C-E**). Noting that miRNA regulators are avidly binding the C5’s intronic regions, we next looked at miRNA motif enrichment within the introns using HOMER (65), which revealed significant enrichment for 14 miRNA motifs (**Supplementary Fig. 3A**). C5 IR does not play a role in gene expression regulation, as revealed by the analysis of fold-changes over time of the genes containing the retained introns (**Supplementary Fig. 3B**), and may thus serve other functions, particularly in the cytoplasm. Focusing on miR-4519, miR-1976, miR-4716-5p, miR-485-5p and miR-4267, the top five miRNAs motifs enriched in the C5 intronic sequences (**Fig. 3F**), we next sought to test whether changes in the C5 PIR over time might relate to changes in gene expression of specific miRNA predicted target genes, as a result of trapping/releasing these miRNAs. To this end, we examined the gene expression profile of their predicted target genes by combining two miRNA target prediction algorithms, TargetScan (66) and miRanda (67). Strikingly we find that the predicted target genes of miR-4519, miR-1976, and miR-4716-5p exhibit a reduction in expression from DIV=7 to DIV=14, which coincides with a reduction of the C5 introns PIR, while the predicted target genes of miR-485-5p and miR-4267 do not exhibit such trend (**Fig. 3G** and **Supplementary Figs. 5,6**). This result indicates that the decrease in IR from DIV=7 to DIV=14 correlates with an increase in miRNA activity, supporting the hypothesis that cytoplasmic retained introns reduce miRNA activity potentially by sequestering them, as previously shown for long non-coding RNA (lncRNA) (68).

**Figure 3.**
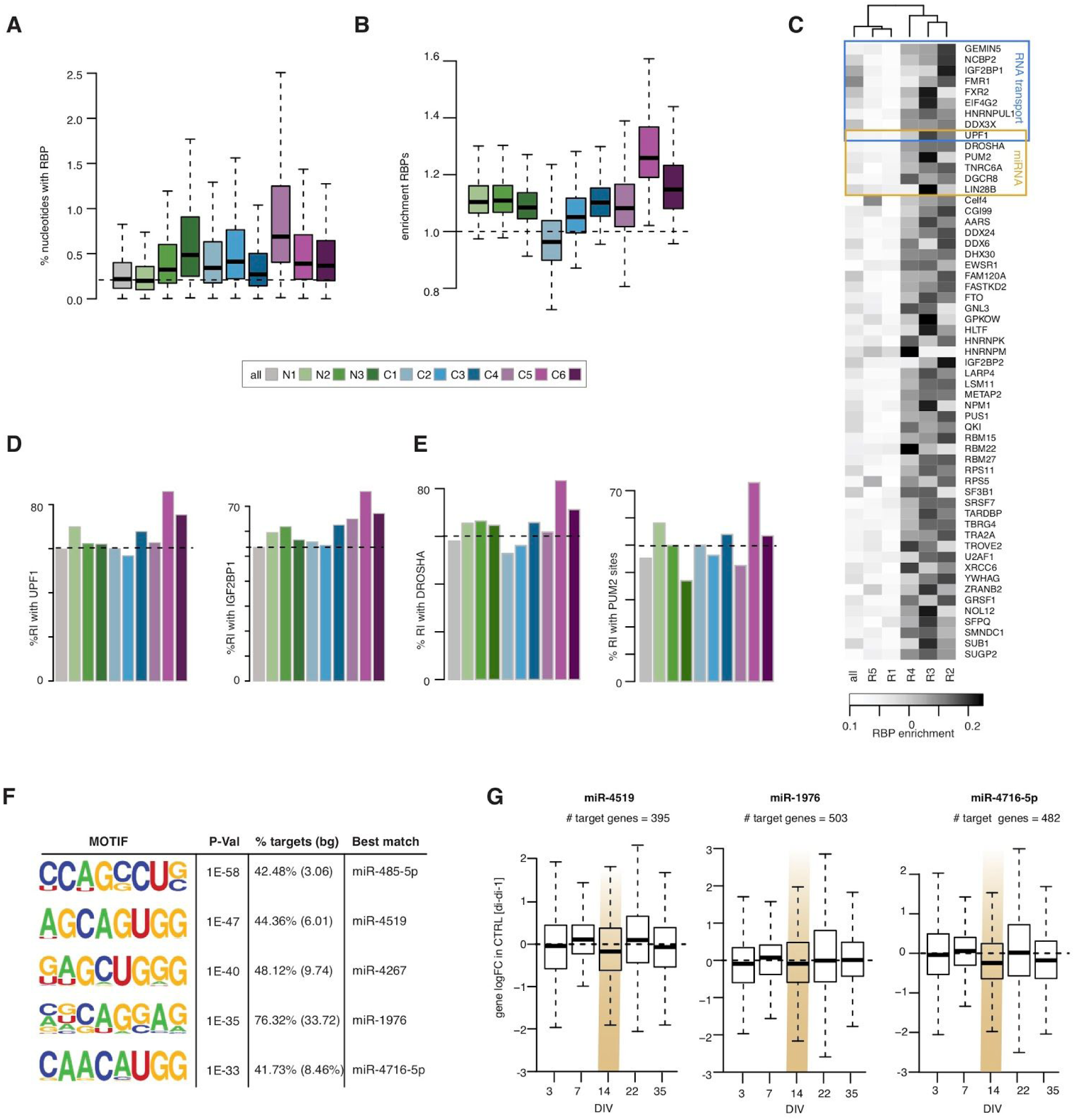
Identification of a cytoplasmic group of retained introns with a high capacity for RBP and miRNA sequestration. **A**. Analysis of the percentage of nucleotides with cross-linking events for 131 RBPs across the entire retained intron for all 9 categories. **B**. Analysis of the enrichment in binding sites for 131 RBPs across the entire retained intron for all 9 categories. Enrichment is obtained by dividing the fraction of retained introns in the category of interest with a CLIP binding with the fraction of retained introns in the complete set of introns (n=61872) with a crosslinking event. **C**. Heatmap of the enrichment score of crosslinking events in both the entire intron and each of the five regulatory regions of the 270 retained introns of the C5 category for 51 RBPs that exhibit a difference in the fraction of cross-linking events of more than 19% in the pool of C5 retained introns compared to the full set of introns. The blue box highlights RBPs involved in RNA transport and the gold box represents those involved in miRNA regulation. **D, E**. Percentage of retained introns with crosslink events for two RBPs involved in RNA transport (UPF1 and IGF2BP1), and two RBPs involved in miRNA regulatory pathway (DROSHA and PUM2). **F**. Five top motifs enriched in the 270 retained introns of the C5 category identified by HOMER (65). **G**. Distributions of the changes in nuclear expression over time of the control samples for the TargetScan (66) and miRDB (78) predicted target genes of miR-4519, miR-1976 and miR-4716-5p. Fold-changes over time obtained by comparing the log2 expression level at time of interest (*d*_*t*_ = {0, 3, 7, 14, 22, 35}) with the expression level at previous stage (*d*_*t*−1_). Gold shaded area indicates the time-point where the largest changes in cytoplasmic IR are observed over time for the control samples. For A,B & G: Data shown as box plots in which the centre line is the median, limits are the interquartile range and whiskers are the minimum and maximum.

### VCP mutation-related transient accumulation of cytoplasmic IRTs correlates with reduced miRNA activity

We previously demonstrated aberrant cytoplasmic IRTs in ALS-related VCP^*mu*^ samples during MN differentiation that exhibit high predicted binding affinity for RBPs (20, 23). We next sought to test whether VCP mutations affect any of the cytoplasmic groups of introns in particular. Examining the two most prominent classes of cytoplasmic IR dynamics during MN differentiation, as captured by right singular vectors of the SVD analysis performed on the cytoplasmic PIR values, confirmed prior findings that VCP mutations leads to exceptionally large IR perturbations at DIV=14 (**Supplementary Figs. 4A, B**) (23). Further comparing the PIR distributions between control and VCP^*mu*^ samples in each of the six groups of cytoplasmic retained introns revealed that VCP mutations specifically impact two classes of events, namely C5, and to a much lesser extent C1 while the other groups remain unchanged (**Fig. 4A** and **Supplementary Fig. 4C**). Most notably VCP-driven changes in cytoplasmic IR are 1) unidirectional i.e. we only detect increases in IR in VCP^*mu*^ samples compared to control samples irrespective of PIR dynamics, and 2) the VCP mutation specifically affects groups of introns in which the PIR exhibits a large decrease from DIV=7 to DIV=14. This is in contrast to those groups of introns where the PIR *increases* from DIV=7 to DIV=14, such as C2 and C6, where we find similar increase in control and VCP^*mu*^ samples (**Supplementary Figs. 4C**). These results suggest that VCP mutations enhance the cytoplasmic stability of IRTs, rather than affecting nuclear export, which would equally impact C1, C2, C5 and C6.

**Figure 4.**
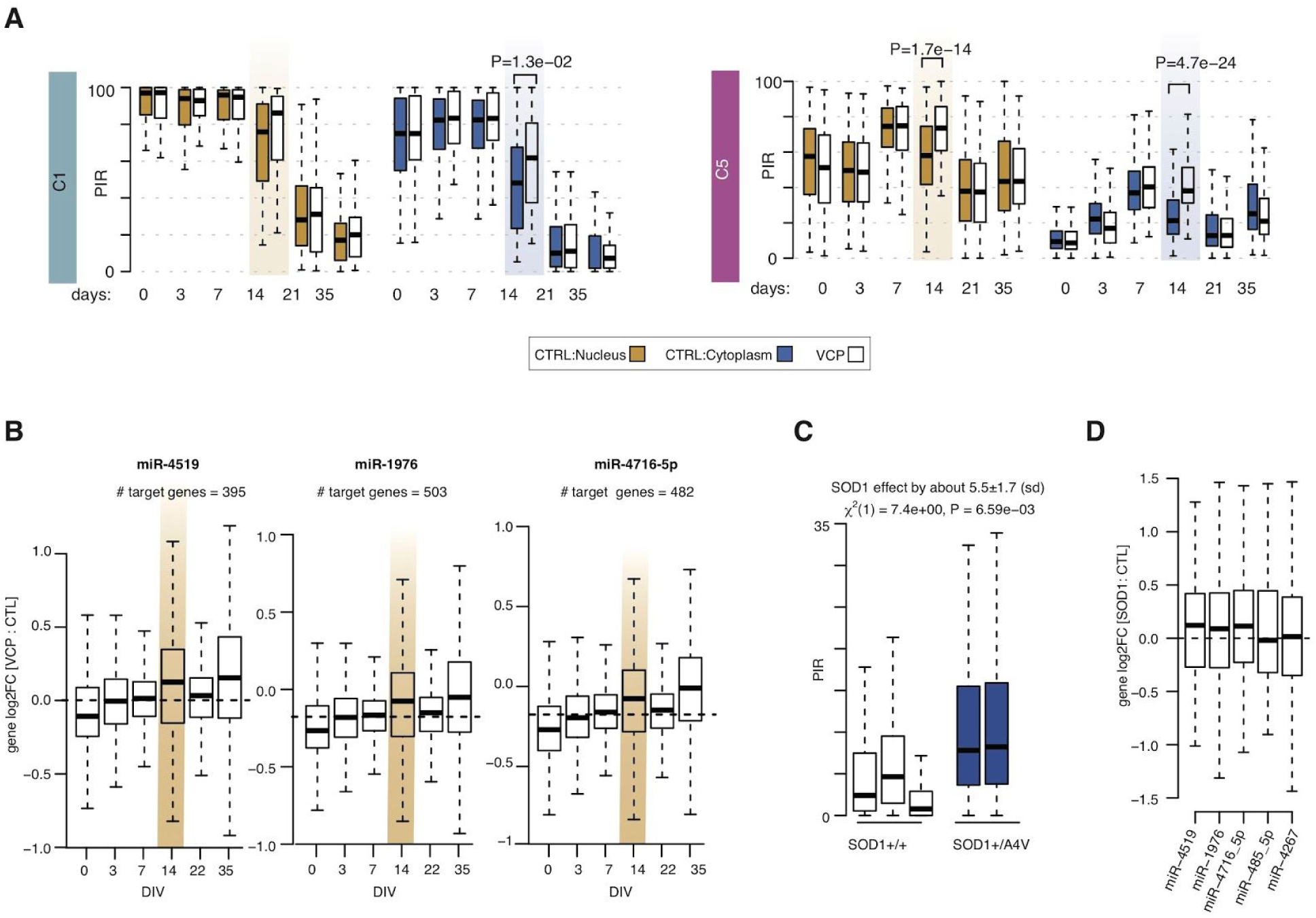
ALS-related transient accumulation in cytoplasmic retained introns correlates with reduced miRNA activity. **A**. Comparison of the distributions of nuclear and cytoplasmic percent intron retention (PIR) between control (*colored* boxes) and VCP^*mu*^ (*white* boxes) samples during MN differentiation for the C1 and C5 groups of cytoplasmic retained introns. P-values obtained with two-sided Welch *t*-test. **B**. Distributions of the changes in nuclear expression between VCP^*mu*^ and control samples at each time-point (*right*) for the TargetScan (66) and miRDB (78) predicted target genes of miR-4519, miR-1976 and miR-4716-5p. Fold-changes obtained by comparing the log2 expression level at each time point (*d*_*t*_ = {0, 3, 7, 14, 22, 35}). Gold shaded area indicates the time-point where the largest changes in cytoplasmic IRTs are observed between the control and VCP mutant samples. **C**. Boxplots displaying the distribution of percentage retention for the 270 introns of the C5 category in control MNs (white box), and SOD1^*mu*^ MNs samples (*left*, blue bar) (79, 80). Mutant samples exhibit a systematically higher proportion of IR compared with controls. Linear mixed effects analysis of the relationship between the PIR for the 270 introns and SOD1 mutation to account for idiosyncratic variation due to patient differences: SOD1 mutation significantly increases IR by about 5.6% ± 3 (standard errors; χ ^2^ (1) = 7.4, P = 6.5E-03). **D**. Distributions of the changes in expression between SOD1^*mu*^ and control samples for the predicted target genes of miR-4519, miR-1976, miR-4716-5p, miR-485-5p and miR-4267. For A-D: Data shown as box plots in which the centre line is the median, limits are the interquartile range and whiskers are the minimum and maximum.

Having found that VCP mutations lead to large perturbations of the C5 group of cytoplasmic IRTs at DIV=14, which we have shown above to associate with decreased activity of specific miRNAs, we next sought to test whether the increase in cytoplasmic IR of the C5 group in VCP mutants correlates with a reduction in miRNA activity by looking at the changes in gene expression of their predicted target genes between VCP and control samples. This analysis revealed that the increase in IR in VCP mutant cultures correlates with a decrease in miR-4519, miR-1976 and miR-4716-5p activities as predicted by the up-regulation of their respective target genes at DIV=14 (**Fig. 4B**). Additionally, these changes are not explained by a change in the expression levels of these miRNAs which are not significantly different between VCP and control cultures at this time point (**Supplementary Figure 4D**). Notably, the predicted activities of miR-485-5p and miR-4267, whose target gene expression profiles did not correlate with IR level in control samples over time, also do not correlate with increase in C5 IR in VCP samples, thus supporting the hypothesis that the activities of the same miRNAs correlate to IR in both VCP and control samples over time (**Supplementary Fig. 6**).

Noting that we have studied a relatively rare form of familial ALS (fALS) caused by gene mutations in VCP (selected as it exhibits the pathological hallmark of TDP-43 nuclear-to-cytoplasmic mislocalisation), we next sought to understand the generalizability of the association between increase in IR in ALS samples and decrease in miRNA activity. To this end, we chose to study one of the most common forms of fALS (SOD1), which in contrast does not exhibit the pathological hallmark of TDP-43 nuclear-to-cytoplasmic mislocalisation. We first looked at the PIR of the C5 group in SOD1 mutant hiPSC-derived MNs, revealing a statistically significant increase in IR in SOD1 (**Fig. 4C**). Next, looking at the changes in gene expression of the miRNA predicted target genes between SOD1 mutant MNs and their isogenic controls, further showed a decrease in miR-4519, miR-1976 and miR-4716-5p predicted activities (**Fig. 4D**). These findings further substantiate the relevance of the correlation between increased IR and decreased miRNA activity. Altogether these findings support the hypothesis that the cytoplasmic pool of C5 introns leads to a reduction in miRNA activity, potentially through direct binding and sequestration, which may have important roles in ALS pathogenesis, and indeed implications for new therapeutic strategies.

## DISCUSSION

Neuronal biology relies on complex regulation of gene expression and mRNA metabolism. Alternative splicing has been shown to play a key role in this process and IR is now recognized as the dominant mode of splicing during MN development (1, 20), including cytoplasmic IR, which we recently showed to affect >100 transcripts during neuronal development (23). Because nuclear IR has been the focus of most previous studies, the regulation and role of cytoplasmic IRTs remain unclear. The objective of this study was twofold: to deepen our understanding of the role(s) of cytoplasmic IR in normal cellular physiology by resolving the spatiotemporal dynamics of IR underlying distinct stages of MN lineage restriction, and to decipher whether specific classes of IRTs become dysregulated in the context of disease by systematically examining the influence of ALS-causing VCP mutations on this process. In order to achieve this we re-analyzed nuclear and cytoplasmic RNA-sequencing data from a time course of patient-specific iPSCs differentiating into spinal MNs.

We first show that nuclear and cytoplasmic IR target distinct classes of mRNA associated with particular dynamics, biological pathways and molecular characteristics. Specifically, we find that the sequences of the retained introns that localise to the cytoplasm are evolutionarily more conserved and exhibit a higher capacity for RBP binding compared to the nuclearly detained introns. This argues against the hypothesis that cytoplasmic intron-containing pre-mRNAs simply ‘leak’ from the nucleus (69), which is also further excluded by polyA selection during library preparation, and suggests that 1) cytoplasmic localisation signals for these IRTs are contained in the intronic sequences, and 2) cytoplasmic IRTs likely serve a biological function that has yet to be discovered.

We next show that MN differentiation exhibits complex IR spatiotemporal dynamics captured by 9 distinct IR programmes, 3 which are nuclearly detained and 6 that localise to the cytoplasm. Given the time and cell compartment specificity of these programmes, they are expected to associate with distinct complex regulation. IR has been previously proposed to be the consequence of globally inefficient splicing (47, 70), that could be linked to several mechanisms including the occupancy of MeCP2 near the splice junction (71), the expression of PRMT5 (7), and relatively weak splice sites (1). Here we find that the 9 groups of introns exhibit similar 5’ and 3’ maximum entropy scores as well as similarly high RBP binding in their exonic regions juxtaposed to the splice sites where the splicing machinery is more likely to form (53, 54) as opposed to the intronic regions. These findings indicate that an overall change in splicing efficiency during MN differentiation is unlikely to be the dominant regulatory factor for most of these IR programmes. Furthermore, IR during MN differentiation does not occur stochastically, but appears to target a specific set of introns in each gene, and thus an additional layer of specific regulation must underlie the regulation of these 9 distinct IR programmes as previously suggested (47). Indeed similar combinations of trans-acting factors are detected across 4 regions juxtaposed to the splices sites among 3 groups of cytoplasmic retained introns -{C4, C5, C6}-suggesting similar regulation. Additionally these three groups of introns exhibit avid RBP binding within their intronic regions juxtaposed to the splice sites when compared to the full set of analysed introns, indicating a potential regulatory mechanism in mRNA metabolism through intronic sequence binding for the {C4, C5, C6} groups of introns. Notably the full list of retained introns for each group together with the regional RBP enrichment is freely accessible as supplementary tables providing a rich resource for researchers across the disciplines of genomics and basic neuroscience.

Although the stable cytoplasmic localisation of intronic sequences in neurons has been recognized since 2013 (11), their role has remained poorly understood. One of the few studies focussing on cytoplasmic IR showed an ‘addressing’ function for intronic RNA sequences in determining their spatial localization within cellular compartments (15). Here we show that the avid RBP binding we previously observed in ALS-related aberrant cytoplasmic retained introns (23) is indeed specifically detected in one IR programme which exhibits a transient increase in the cytoplasm during MN differentiation, namely the C5 group. Notably this group of cytoplasmic intron, which is the most impacted by VCP mutations, exhibits the same PIR dynamics of the group of introns we previously showed to be impacted by VCP mutations using whole-cell RNA-sequencing (20). This suggests that VCP mutations specifically affect the cytoplasmic stability of IRTs rather than leading to a reduction in splicing efficiency. The absence of correlation between the C5 PIR level and the gene expression dynamics during MN differentiation raises the possibility of new roles for intronic RNA sequences beyond a function in gene expression regulation, particularly in the cytoplasm. As previously proposed, cytoplasmic retained introns may act as RNA regulators in the homeostatic control of RBP localisation during development and disease (23), which may in turn lead to loss of function. For example some splicing factors that avidly bind the C5 intronic sequences may remain sequestered upon their nuclear export and cytoplasmic localisation, contributing to a transient reduction in splicing efficiency during MN differentiation. Another intriguing molecular characteristic of the C5 group of introns is the enrichment for several miRNA motifs across the full length of the intron, predicted activity for which negatively correlates with the PIR level of this intron group during MN differentiation. Thus the presence of long intronic sequences in the cytoplasm of neuronal cells may serve as a regulatory mechanism for miRNA functionality through their sequestration and downstream up-regulation of their target genes. Indeed previous studies speculated that stable intron-derived RNA sequences (sisRNA) (72) act as molecular sinks to sequester miRNA (72) and/or RBPs (73) leading to reduction in their activities and future studies will test whether sisRNA are derived from the cytoplasmic intronic sequences of the C5 group.

We previously demonstrated aberrant cytoplasmic IR in ALS-related VCP^*mu*^ samples during MN differentiation (23). Here we show that VCP mutations lead to an aberrant PIR increase specifically of the C5 group of introns. Furthermore we show that the aberrant increase in C5 PIR level at DIV=14 in ALS mutant cells correlates with a decrease in the predicted activities of miR-1976, miR-4519 and miR-4716-5p, motifs of which are enriched in the C5 intronic sequences. These findings were further generalized to SOD1-related ALS hiPSC-derived mutant MNs, supporting the hypothesis of a functional depletion of specific miRNAs as a result of cytoplasmic intronic sequences-mediated sequestration in ALS cells. Notably a reduction in miR-1976 activity, motifs of which are detected in 76% of the C5 intronic regions, is expected to occur in some sporadic ALS patients due to a mutation (rs17162257) in its enhancer (74). Furthermore, several miRNAs, and their target genes, are recognized to be involved in the occurrence and pathophysiology of neurodegenerative diseases including ALS (75–77). Thus, here we propose that a group of intronic sequences which accumulate in the cytoplasm of VCP mutant cells, as previously shown (23), act as molecular sponges for miRNA, thus resulting in elevated expression of their target genes. Future work will be required to demonstrate the direct role of these cytoplasmic intronic sequences in regulating miRNA activity through sequestration.

In conclusion we propose that cytoplasmic retained introns function as RNA regulators in the homeostatic control of RBP localisation and miRNA activity during MN development and disease, which has potential implications for ALS pathogenesis and the development of therapies for this devastating and incurable disease.

## FIGURES AND TABLES

**Supplementary Figure 1.**
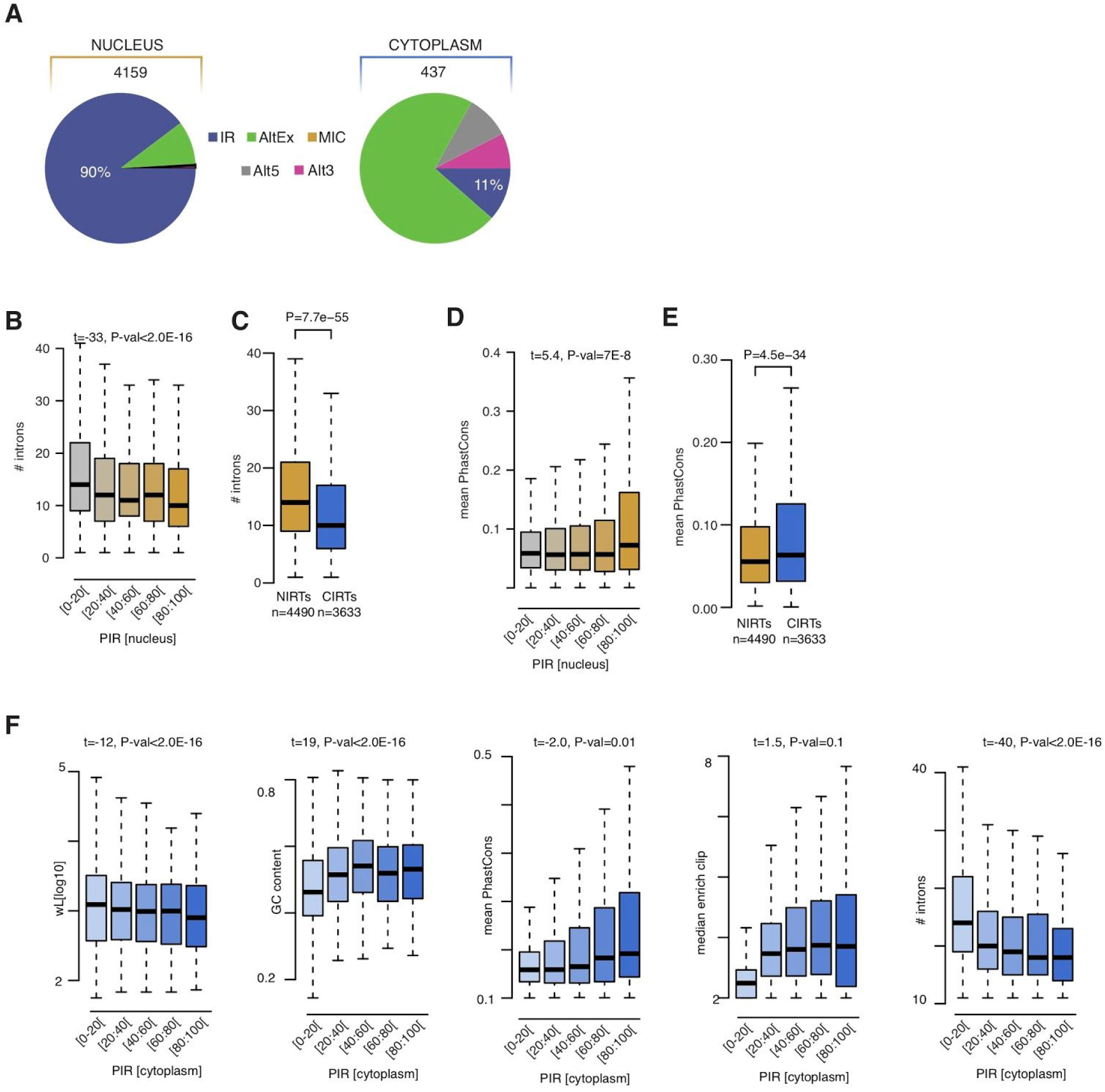
**A**. Pie charts representing proportions of skipped splicing events at defined stages during motor neurogenesis in nuclear (*left*) and cytoplasmic (*right*) fractions. Total number of events are indicated above the charts. Intron retention (IR); alternative exon (AltEx); microexons (MIC); alternative 5′ and 3′ UTR (Alt5 and Alt3). **B, D**. Analysis of the relationship between the percent intron retention (PIR) in the nucleus and the number of introns per gene (**A**), and the retained intron average conservation scores (**D**). Retained introns are grouped in five categories of increasing level of retention in the nucleus as indicated on the *x*-axis. Data shown as in Fig. 1D. *P*-values obtained from analysis of variance comparing the full model of the logit of maximum IR across all nuclear samples according to the five characteristics with the reduced model removing the characteristic of interest. **C, E**. Comparison of number of introns per gene and the conservation scores between nuclear and cytoplasmic retained introns. Nuclear retained introns are defined as those exhibiting >20% IR in nuclear fraction and <5% IR in cytoplasmic fraction. Cytoplasmic retained introns are defined as those exhibiting >20% IR in nuclear fraction and >15% IR in cytoplasmic fraction. Data shown as in Fig. 1D. P-values obtained from Mann-Withney test. **F**. Analysis of the relationship between the percent intron retention (PIR) in the cytoplasm and the intron length, the GC content in %, the number of introns per gene, the retained intron average conservation scores and the median enrichment for RBP binding site compared to the non-retained introns of the same gene. Retained introns are grouped in five categories of increasing level of retention in the cytoplasm as indicated on the *x*-axis. Data shown as box plots in which the centre line is the median, limits are the interquartile range and whiskers are the minimum and maximum.

**Supplementary Figure 2.**
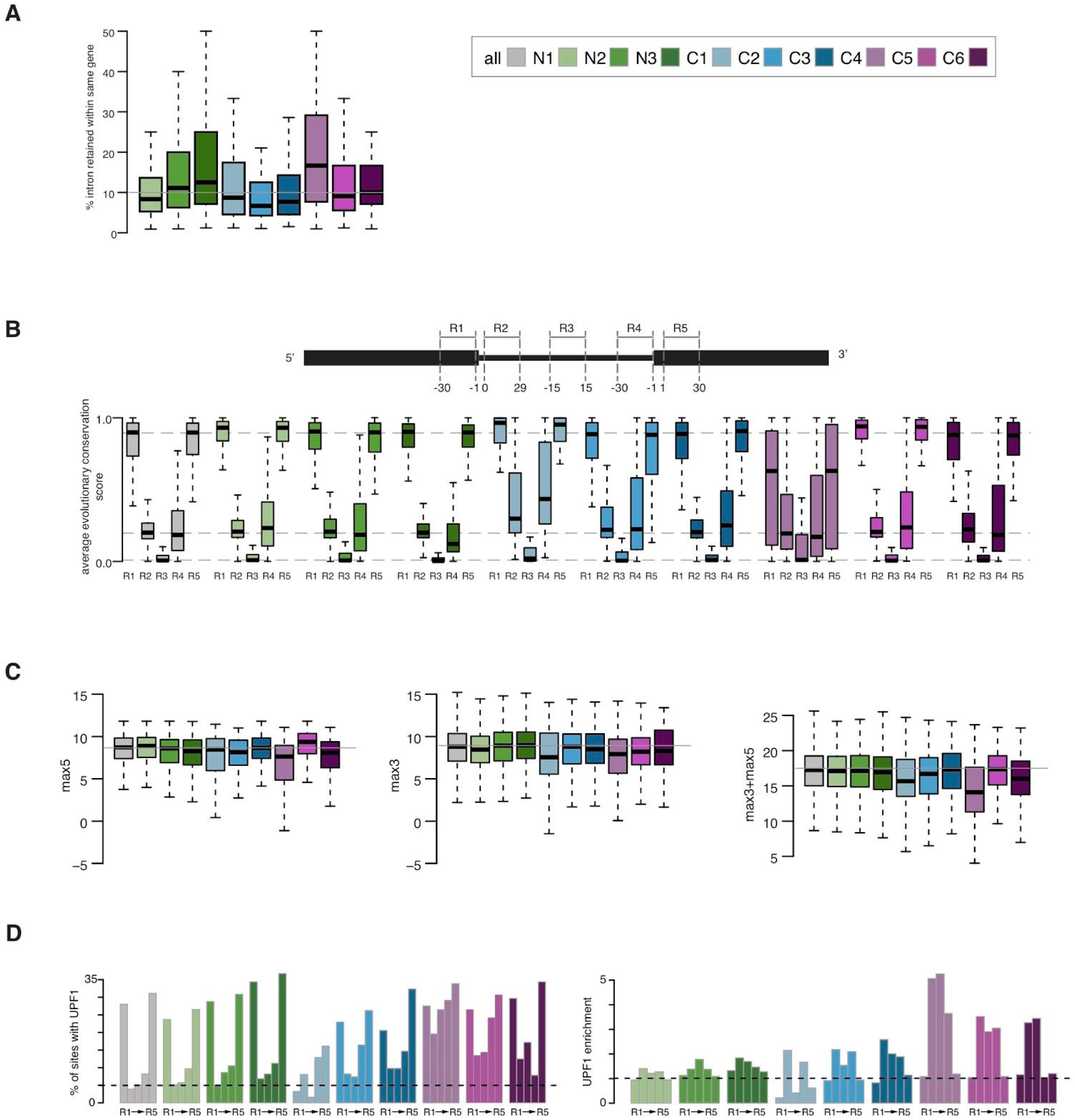
**A**. Percentage of retained introns per gene for the genes targeted by intron retention in each group. **B**. Distribution of the average evolutionary sequence conservation scores in the in the last 30nts of the upstream exon (R1), the first 30nts of 5’ intron region (R2), the 30nts in the middle of the intron (R3), the last 30nts of 3’ intron region (R4), and the first 30nts of downstream exon (R5) for 9 categories of introns for the 9 categories of introns. **C**. Distribution of the maximum entropy scores for 9-bp 5′ splice sites and 23-bp 3′ splice sites for the 9 categories of intron as obtained from MaxEntScan (38). **D**. Percentage of introns with UPF1 regional cross-linking events (*left*) and UPF1 regional cross-linking enrichment (*right*) for each splicing regulatory regions R1, R2, R3, R4 and R5 in each group of introns. Dashed lines indicate the average percentage of all 61872 analysed introns with a CLIP binding (*left*) and the one-fold enrichment (*right*) in the intronic regulatory regions (R2, R3, R4).

**Supplementary Figure 3.**
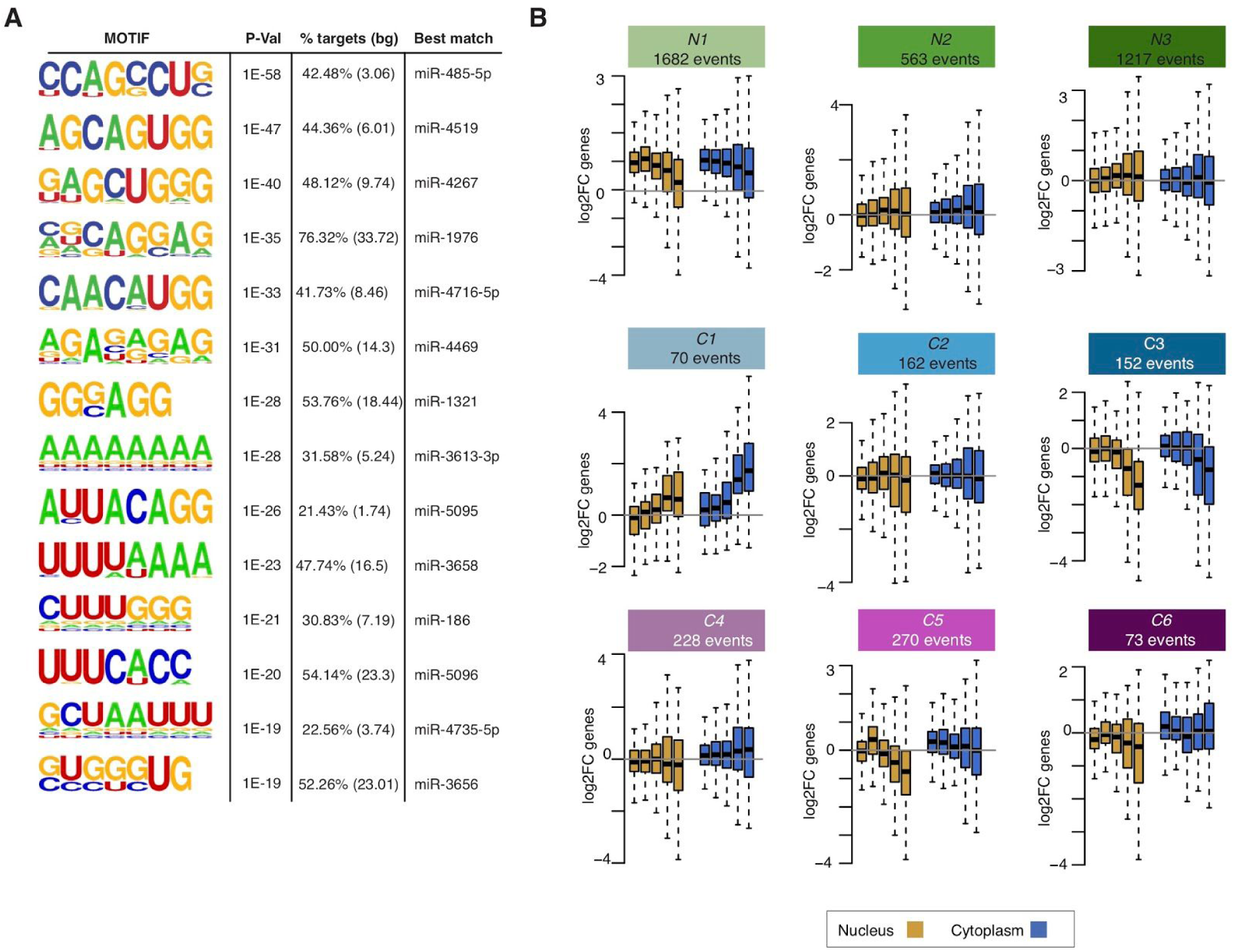
**A**. 14 motifs enriched in the 270 retained introns of the C5 category identified by HOMER (65). **B**. Changes in gene expression over time inthe nucleus (*gold* boxes) and cytoplasm (*blue* boxes) for groups of genes containing the 9 different categories of retained introns. Fold-changes obtained by comparing the log2 expression level at time of interest (*d*_*i*_ = {3, 7, 14, 22, 35}) with the expression level at iPSC stage (*d*_0_). Data shown as box plots in which the centre line is the median, limits are the interquartile range and whiskers are the minimum and maximum.

**Supplementary Figure 4.**
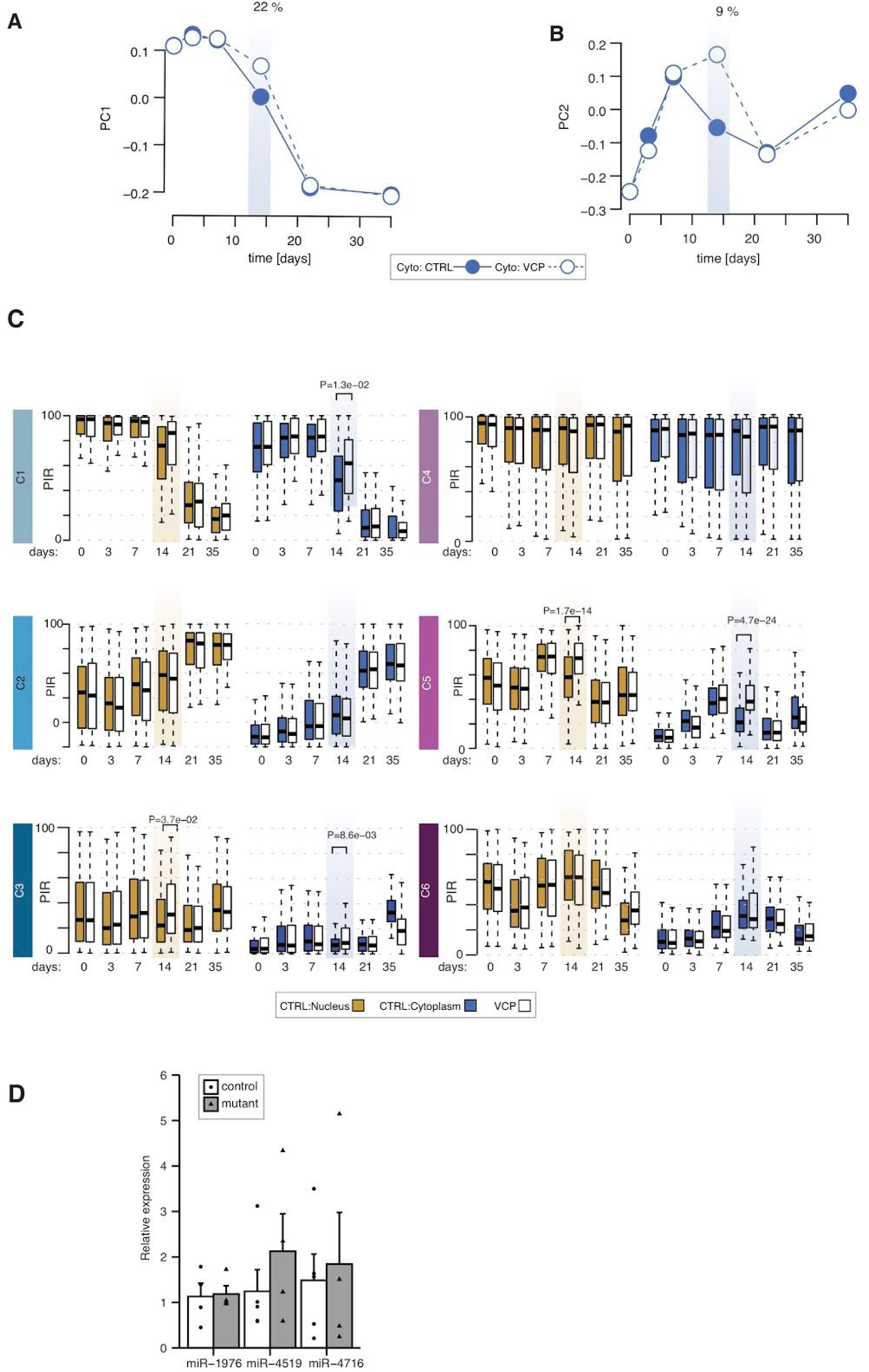
**A, B**. Singular value decomposition analysis of the PIR cytoplasmic values of 94,457 introns in n = 48 cytoplasmic samples. Line plots showing the PIR profiles of the first two singular vectors *v*_1_ and *v*_2_, capturing 22% and 9% of the variance in PIR respectively. Filled and empty data points indicate PIR values for the control and VCP^*mu*^ samples. **C**. Comparison of the distributions of nuclear and cytoplasmic PIR between control (*colored* boxes) and VCP^*mu*^ (*white* boxes) samples during MN differentiation for the 6 groups of cytoplasmic retained introns. P-values obtained with two-sided Welch *t*-test. Data shown as box plots in which the centre line is the median, limits are the interquartile range and whiskers are the minimum and maximum. **D**. MiRNA expression in “patterned” precursor motor neuron cells (DIV=14) – Relative expression of miR-1976, miR-4519 and miR-4716 in control (white) and mutant (gray) cells lines. Datapoints depicted as black circles in the controls and black triangles in mutant cells.

**Supplementary Figure 5.**
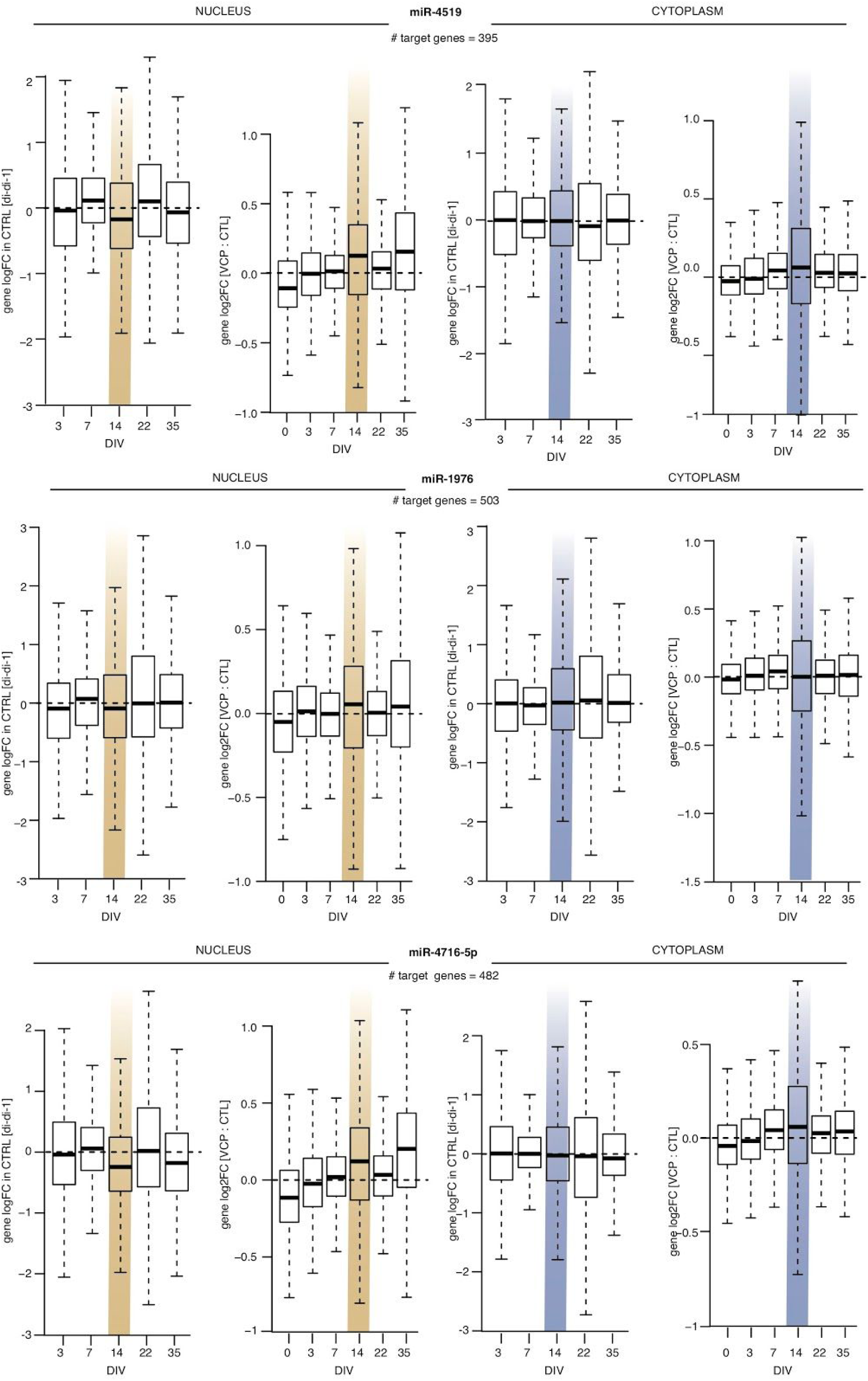
Distributions of the changes in nuclear and cytoplasmic expression over time of the control samples (*left*) and between VCP^*mu*^ and control samples at each time-point (*right*) for the 395 TargetScan (66) and miRDB (78) predicted target genes of miR-4519, miR-1976, and miR-4716-5p. Fold-changes over time obtained by comparing the log2 expression level at the time of interest (*d*_*t*_ = {0, 3, 7, 14, 22, 35}) with the expression level at previous stage (*d*_*t*−1_). Gold shaded area indicates the time-point where the largest changes in cytoplasmic IR are observed either over time for the control samples or between control and mutant samples.

**Supplementary Figure 6.**
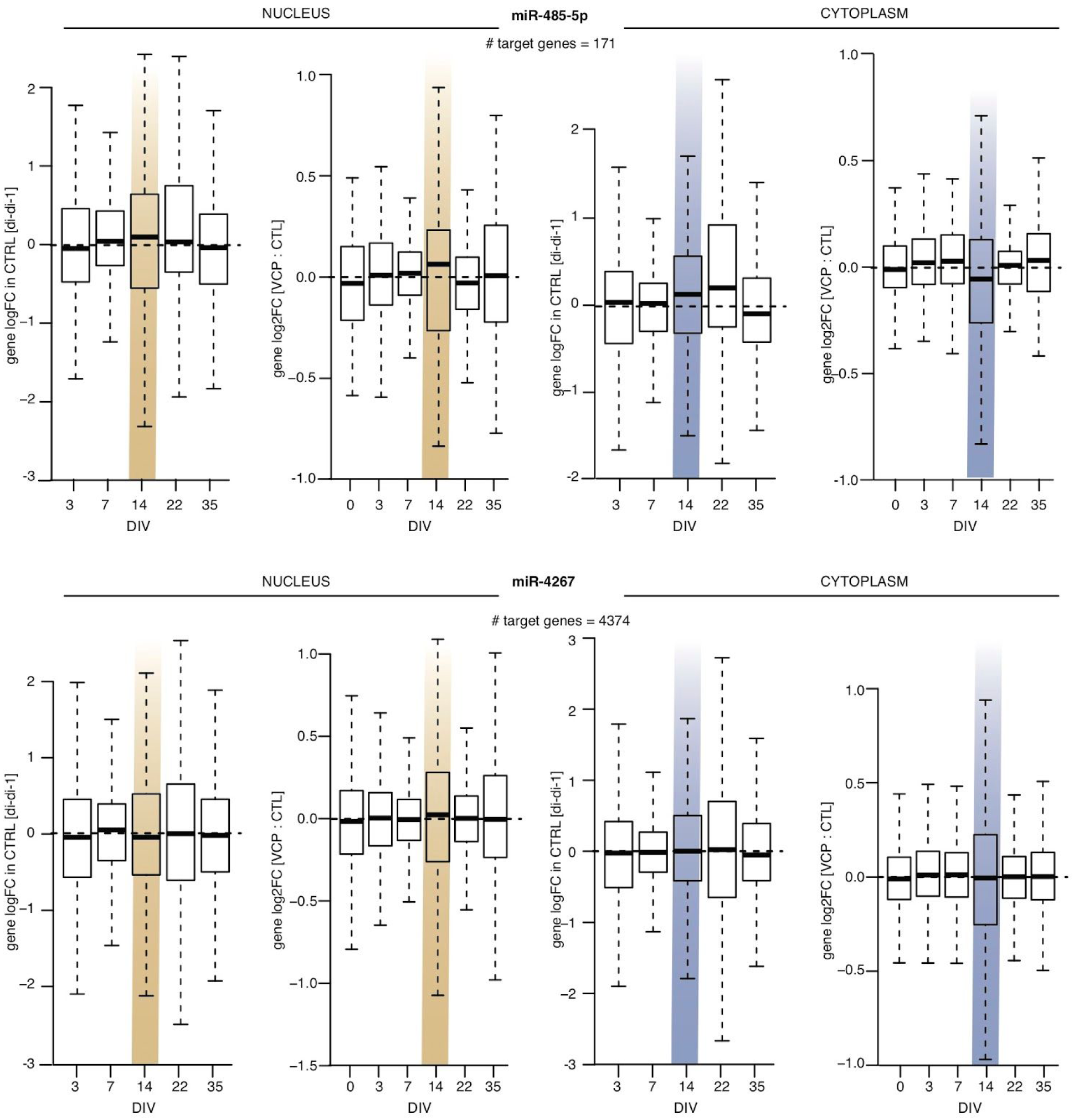
Same details and format as Supplementary Figure 5 but for miR-485-5p and miR-4267.

## SUPPLEMENTARY TABLES

**1-18** can be accessed here.

**Table S1** | Description of the iPSC lines and RNA sequencing samples used in this study.

**Table S2** | List of the 4490 nuclearly detained retained introns reported on Figs. 1F,H,J.

**Table S3** | List of the 3633 cytoplasmic retained introns reported on Figs. 1F,H,J.

**Tables S4-S12** | Lists of the 9 groups of retained introns (N1, N2 N3, C1, C2, C3, C4, C5, C6) associated with the distinct spatiotemporal dynamics during MN differentiation reported on Fig. 2A.

**Tables S12-S17** | Frequency and enrichment in 133 RBPs crosslink events in 5 defined regions of interest (R1, R2, R3, R4, and R5).

## AUTHOR CONTRIBUTIONS

Conceptualization, R.L., R.P.; Formal Analysis, R.L.; Investigation, R.L., M.P-H., H.C., J.N., G.E.T.; Writing – Original Draft, M.P-H., R.L., R.P.; Writing – Review & Editing, M.P-H., H.C., J.N., G.E.T., R.L., R.P; Resources, R.L., R.P.; Visualization, R.L.; Funding Acquisition, R.L., R.P.; Supervision, R.L, R.P.

## ACKNOWLEDGMENTS

The authors wish to thank the patients for fibroblast donations. We also thank Anob M Chakrabarti for sharing BED files of aligned CLIP data. We are grateful for the help and support provided by the Scientific Computing section and the DNA Sequencing section of Research Support Division at OIST. This work was supported by the Idiap Research Institute and by the Francis Crick Institute which receives its core funding from Cancer Research UK (FC010110), the UK Medical Research Council (FC010110), and the Wellcome Trust (FC010110). D.M.T. is supported by a Newton-Mosharafa scholarship. R.P. holds an MRC Senior Clinical Fellowship [MR/S006591/1].

